# Conservation of the structural and functional architecture of encapsulated ferritins in bacteria and archaea

**DOI:** 10.1101/431494

**Authors:** Didi He, Cecilia Piergentili, Jennifer Ross, Emma Tarrant, Laura R. Tuck, C. Logan Mackay, Zak McIver, Kevin J. Waldron, Jon Marles-Wright, David J. Clarke

## Abstract

Iron is an essential element for many biological processes; however, due to its high reactivity iron can also be very toxic, producing reactive oxygen species through Fenton chemistry. Ferritins protect the cell from oxidative stress by catalytically converting Fe(II) into less toxic Fe(III) and storing the resulting iron minerals within their core. Encapsulated ferritins (EncFtn) are a sub-family of ferritin-like proteins, which are widely distributed in all bacterial and archaeal phyla. We recently characterised the *Rhodospirillum rubrum* EncFtn, showing that although enzymatically active, due to its open structure it requires the association with an encapsulin nanocage in order to act as an iron store. Given the wide distribution of the EncFtn family in organisms with diverse environmental niches, a question arises as to whether the structure and catalytic activity is conserved across the family. Here we structurally characterise two EncFtn members from the halophile *Haliangium ochraceum* and the thermophile *Pyrococcus furiosus*, which show the same distinct annular decamer topology observed in *R. rubrum* EncFtn, with the ferroxidase centre (FOC) formed between one of the dimer interfaces. Solution and native mass spectrometr analyses show that the stability of the protein quaternary structure differs between EncFtn proteins from different species. The catalytic role of the EncFtn proteins was confirmed by biochemical assays, and we show that Zn(II) ions inhibit the ferroxidase activity of the EncFtn proteins to varying degrees. Our results represent a further step in the characterisation of the recently discovered EncFtn ferritin-like sub-family, indicating a common structural organisation and catalytic activity, despite diverse host environments.

---

The encapsulated ferritins (**EncFtn**) are recently described members of the ferritin superfamily (1–3). These proteins are sequestered within encapsulin nanocompartments, and the two proteins act in concert to provide an iron storage system with a much greater capacity than the classical ferritins and DNA-binding Protein from Starved cells (DPS) nanocages (3, 4). Genes encoding encapsulin associated ferritins have been identified in a wide range of bacterial and archaeal species that inhabit diverse ecological niches from ponds, streams (*Rhodospirillum rubrum)*, and coastal seas (*Haliangium ochraceum*), to abyssal ocean vents (*Pyrococcus furiosus)* (5).

The *R. rubrum* EncFtn was the first protein in its family to be structurally and biochemically characterised; it has a fold with two antiparallel alpha-helices that adopts an annular-decamer quaternary structure, assembled as a pentamer of dimers (1). The subunits dimerise through metal-mediated contacts to reconstitute the four-helix ferritin fold with a functional ferroxidase centre (6, 7). While the EncFtn protein displays ferroxidase activity, it is not capable of storing iron in the same way as other ferritins due to its open architecture and the lack of an enclosed cavity for iron mineralisation (8, 9); therefore, EncFtn family proteins must be localised to the interior of an encapsulin nanocage for efficient iron storage (1, 3, 5). Localisation to encapsulin nanocages is mediated by a short encapsulation sequence, which is usually appended to the C-terminus of the EncFtn protein chain (1, 3, 5, 10); addition of this sequence to heterologous proteins is sufficient to direct them to the lumen of encapsulins (11, 12).

The encapsulin protein family is structurally related to the HK97 bacteriophage shell protein and they form icosahedral nanocages of between 25 and 35 nm in diameter. Pores are formed at the icosahedral symmetry axes between subunits to allow substrate access to encapsulated enzyme cargoes (3, 5, 10, 13). The *R. rubrum* encapsulin was shown to associate with over 2000 iron ions per capsid, while being catalytically inactive; and, in concert with EncFtn, capable of mineralising and storing more than four times as much iron as a classical ferritin nanocage (1). Given the wide distribution of encapsulin proteins across bacterial and archaeal phyla, and the conservation of the EncFtn proteins in 20 % of all phyla with encapsulin genes, we investigated the structure and activity of EncFtn proteins from the halophilic Proteobacterium *H. ochraceum* and the EncFtn-fusion from the Euryarchaeota *P. furiosus*, to determine whether the structural and functional organisation seen in the *R. rubrum* EncFtn is conserved within its phylum and between domains and environmental niches.

Here we report the structural and biochemical characterisation of members of the EncFtn family from *H. ochraceum* and *P. furiosus*. Both proteins adopt similar topology to the *R. rubrum* EncFtn and assemble as annular decamers formed from pentamers of dimers, with ferroxidase centres located between one of the dimer interfaces. The ferroxidase activities of the proteins are comparable, and they show similar levels of inhibition by the competing metal ion zinc. These data show that the structural and biochemical features of encapsulated ferritins are conserved across different environmental niches and phyla with particular adaptations for thermal stability in thermophilic microorganisms.

## Results

### Purification of encapsulated ferritins from H. ochraceum and P. furiosus

The encapsulated ferritins from *H. ochraceum* and *P. furiosus* share 18 % amino acid sequence identity with each other, and 58 % and 22 % with the *R. rubrum* EncFtn (respectively referred to as Hoch_EncFtn, Pfc_EncFtn, and Rru_EncFtn herein). Residues shown to be in the ferroxidase centre of the Rru_EncFtn protein are strictly conserved in the Hoch_EncFtn and Pfc_EncFtn proteins, with secondary metal binding sites conserved in Hoch_EncFtn, but not in Pfc_EncFtn (Figure S1).

To explore the structure and function of encapsulated ferritins from different species, we produced examples of this protein family from *H. ochraceum* and *P. furiosus* as truncated variants, lacking the C-terminal encapsulin localisation sequence, by heterologous expression in *Escherichia coli*. The *H. ochraceum* EncFtn was produced as a C-terminal hexa-histidine tagged variant, C-terminal StrepII-tagged variant, and an untagged variant comprising residues 1–98 of the native polypeptide. In *P. furiosus* the encapsulated ferritin forms a single contiguous polypeptide with the encapsulin shell protein; therefore, a truncated version with residues 1–99 encompassing just the EncFtn domain was produced as both a C-terminal hexahistidine tagged variant, a C-terminal StrepII-tagged variant, and an untagged variant. Following purification, all recombinant proteins were analysed by LC-MS to verify their molecular mass. Average neutral masses were obtained for each homologue by liquid chromatography mass spectrometry (Table S1). These all agree with the predicted masses of each protein within 1–2 Da, with some of the protein variants lacking full processing of the initiating methionine.

To interrogate the behaviour of the EncFtn homologues in solution, both the hexahistidine and StrepII tagged variants were subjected to S200 size-exclusion chromatography, calibrated with standards of known molecular weight, and followed by SDS-PAGE analysis (Fig. 1 and Fig S2). Hoch_His elutes as three peaks of increasing area: a small peak at 70 ml consistent with the decamer size of the *Rhodospirillum rubrum* EncFtn; a peak at 81 ml consistent with a dimer; and one at 87 ml corresponding to the monomer fraction. Pfc_His eluted as a single peak at 65 ml, which has a slightly larger apparent size than the decamer fraction of Rru_EncFtn (Fig. 1A). When subjected to SDS-PAGE the Hoch_His peak fractions were partially resistant to SDS and heat-induced denaturation, presenting bands at the approximate molecular weight of a monomer and dimer species (Fig. S2A); the Pfc_His peak fractions were almost fully resistant to SDS and heat denaturation with the majority of protein appearing as a band with an apparent molecular weight of the decamer, with a small proportion of monomer (Fig. S2B). These observations are in accord with the behaviour of the Rru_His variant (Fig S2C). The StrepII-tagged variants of the proteins behave slightly differently in solution (Fig 1B), Hoch_Strep elutes as a monomer; while the Pfc_Strep has a major decamer peak with an additional peak consistent with higher-order aggregation, such as dimers of decamers. The Rru_Strep protein elutes primarily as a decamer. Their appearance on SDS-PAGE gels is almost identical to the His-variants, with the appearance of some dimer in the Pfc_Strep fractions (Fig S2C/D/E).

**Figure 1.**
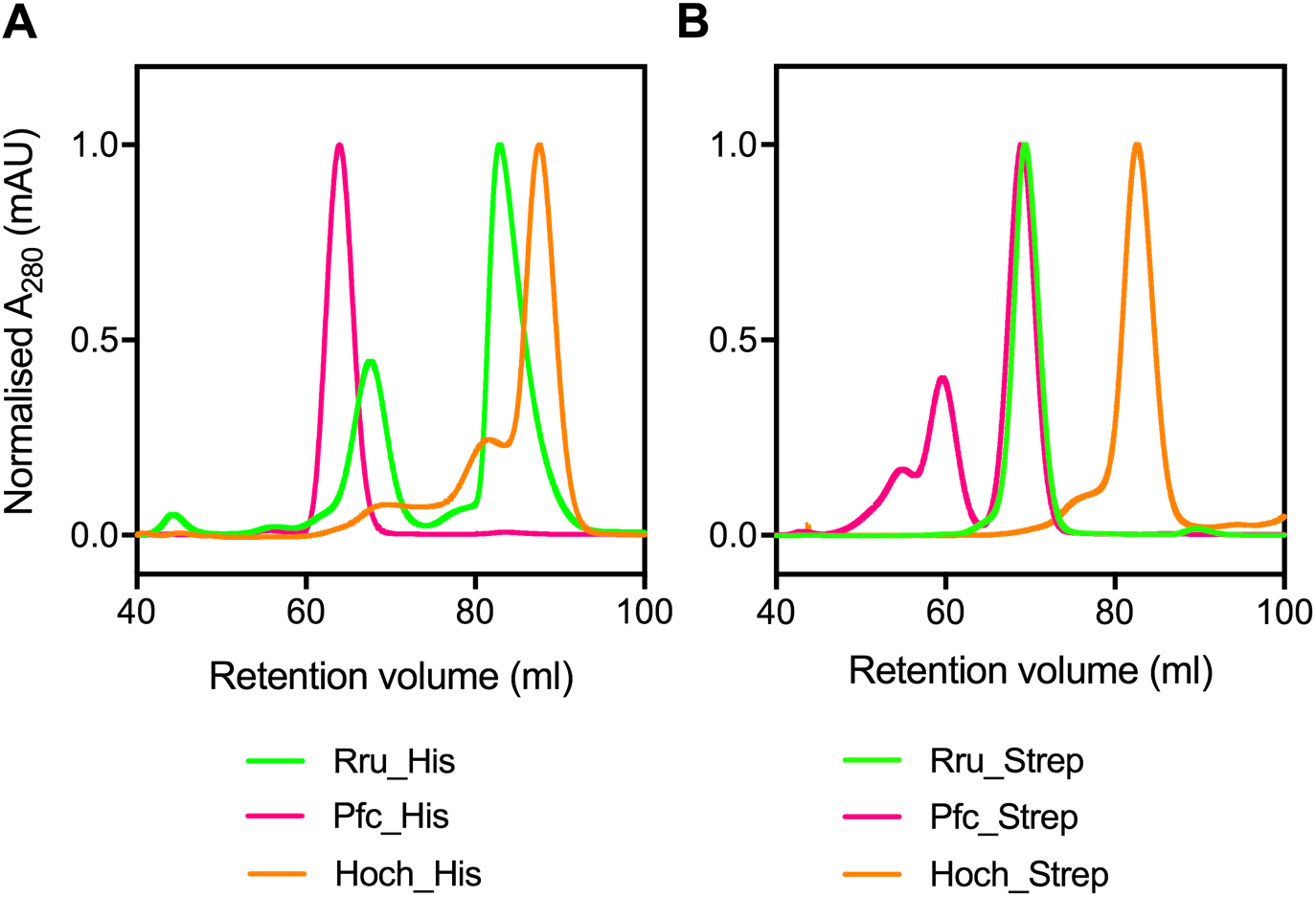
Purification of recombinant *H. ochraceum* and *P. furiosus* EncFtn proteins. *A*, Recombinant hexahistidine-tagged Hoch_His, Pfc_His and Rru_His proteins were purified 94 by nickel NTA-affinity chromatography and subjected to analytical size-exclusion 95 chromatography using a Superdex 200 16/60 column (GE Healthcare) equilibrated with 50 96 mM Tris-HCl, pH 8.0, 150 mM NaCl. The peaks at 70 ml correspond to an estimated 97 molecular weight of 130 kDa when compared to calibration standards, consistent with a 98 decameric species of the EncFtn proteins. The Hoch3836_1–98_ peak at 80 ml corresponds to 99 the 26 kDa dimer and the peaks near 85 ml correspond to the 13 kDa monomer compared to the standards. *B*, Recombinant Strep-tagged Hoch_Strep, Pfc_Strep, and Rru_Strep proteins were purified by Strep-Trap column HP (GE Healthcare). Purified samples were applied to a Superdex 200 16/60 column (GE Healthcare) equilibrated with 50 mM Tris-HCl, pH 8.0, 150 mM NaCl in order to observe oligomerisation state in solution. Rru_Strep elutes primarily at ~ 70 ml, corresponding to the decamer size. Pfc_Strep peaks can be found around 60 ml (~ 20-mer) and at 70 ml (10-mer), whereas Hoch_Strep elutes at ~ 83 ml (1-mer). DOI: 10.6084/m9.figshare.7105907

### Native Mass spectrometry analysis

In order to further understand the differences seen in the solution-phase oligomerisation states of the EncFtn homologues, native mass spectrometry was performed on the hexa-histidine tagged variant of each homologue. As previously reported (1), the decameric assembly of iron-bound Rru_EncFtn can be successfully detected using this technique. Under native MS conditions (at pH 8.0), Rru_EncFtn displays a narrow charge state distribution consistent with the 22+ to 25+ charge state of the protein decamer (Fig. 2A, pink circles). In addition to the decamer, a minor species is observed which is consistent with the iron-free Rru_EncFtn monomer (+6 and +7) (Fig. 2A, blue circles).

**Figure 2.**
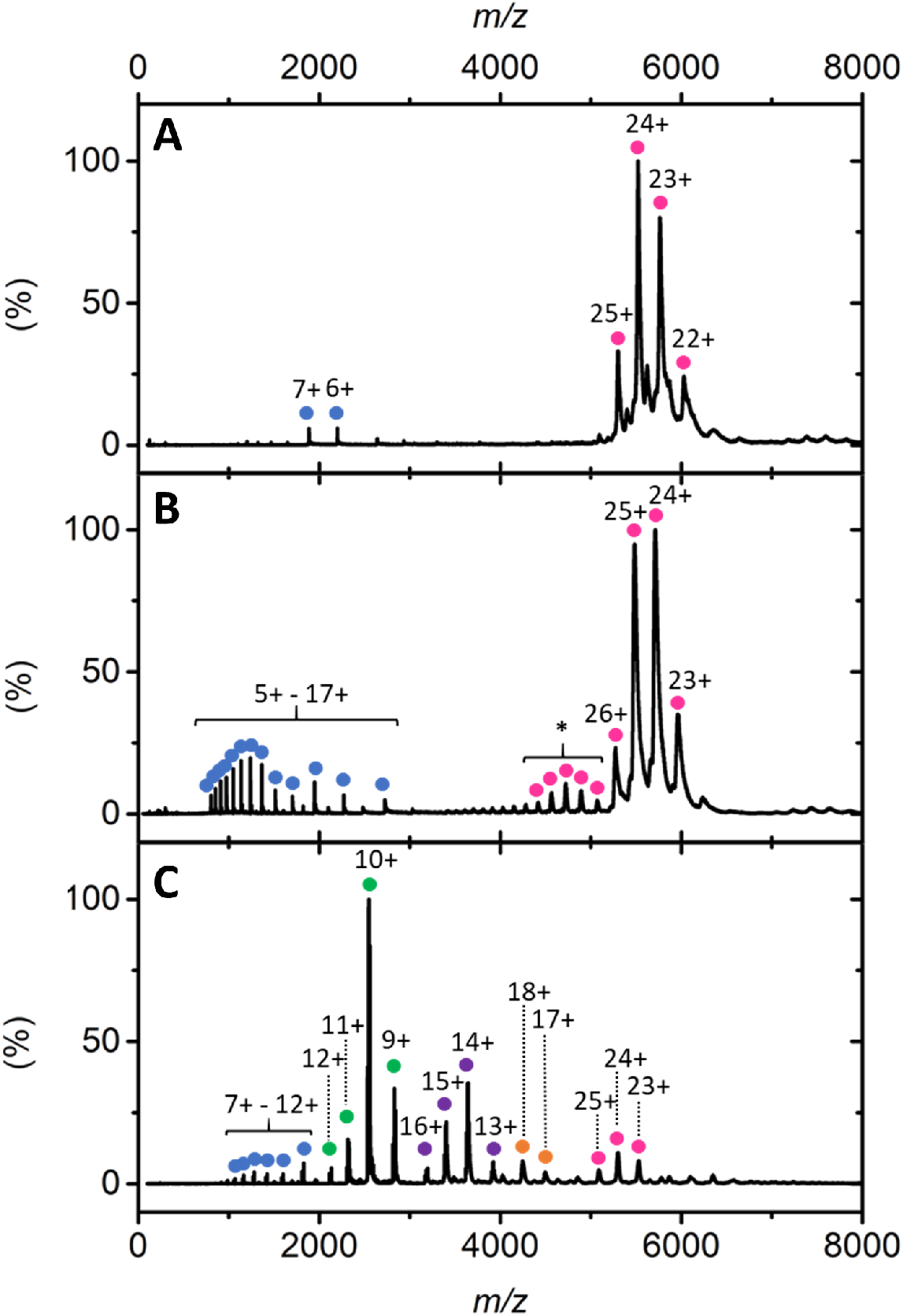
Native mass spectrometry of encapsulated ferritin homologues. Native nanoelectrospray ionization (nESI) mass spectrometry of encapsulated ferritin homologues acquired in 100 mM ammonium acetate (pH 8.0). *A*: nESI spectrum of Rru_EncFtn consistent with decameric assembly. The decameric charge state distribution is represented with pink circles and peaks correspond to 22+ to 25+ charge states. Two minor monomer charge states (blue circles, 6+ and 7+ charge states) are also observed. *B:* Native spectrum of Pfc_EncFtn. Decamer charge states (23+ to 31+) are highlighted with pink circles and monomer charge states (5+ to 17+) are highlighted by blue circles. * denotes the extended charge state observed. *C:* nESI spectrum of Hoch_EncFtn with gas phase oligomerisation stressed with coloured circles. Monomer shown as blue; dimer as green; tetramer as purple; hexamer as yellow and decamer as pink.

In a similar manner to Rru_EncFtn, native MS analysis of Pfc_EncFtn demonstrates charge state distributions consistent with a decameric assembly (+23 to +26) (Fig. 2B, pink circles). Interestingly, the decameric charge state distribution is elongated (Fig. 2B, *) and low abundance (27+ to 31+) charge states are also observed. The elongated charge state distribution is only observed in Pfc_EncFtn, and we attribute its presence to the ability of the solvent exposed affinity tag to readily protonate in solution. Similar to the observations from Rru_EncFtn, a charge state distribution consistent with iron-free Pfc_EncFtn monomer (+5 to +17) is also observed (Fig. 2B, blue circles).

In contrast, native MS analysis of Hoch_EncFtn does not reveal a major decamer species. Instead a series of oligomerisation states are observed, the major species is a dimer (+9 to +12) (Figure 2C, green circles); in addition tetramer (+13 to +16), hexamer (17+ and 18+) and decamer (23+ to 25+), and a small amount of monomer (+7 to +12) gas phase oligomerisation states are all clearly observed (Figure 2C, purple circles, orange circles, pink circles, and blue circles respectively). This observation is consistent with the multiple broad peaks obtained during size-exclusion chromatography (Fig. 1). These data suggest that, under our experimental conditions, multiple oligomeric assemblies of Hoch protein exist in both the solution and gas phase. The observation of even-numbered oligomerisation states (i.e dimer, tetramer and hexamer) suggests that one of the two dimer interfaces is partially unstable, and the protein exists as an equilibrium of dimers and higher order multiples of dimers.

### Crystal structures of Hoch and Pfc Encapsulated ferritins

To explore the structure of the EncFtn homologues, crystals of the Hoch_EncFtn and Pfc_EncFtn were produced by standard crystallisation screening methods. Diffraction data were collected at Diamond Light Source and the crystal structures of the Hoch_EncFtn and Pfc_EncFtn proteins were determined by molecular replacement using a decamer of the Rru_EncFtn (PDB ID: 5DA5) as the search model. Data collection and refinement statistics are shown in Table 1. The structure of Hoch_EncFtn was refined at 2.06 Å resolution and contained a decamer in the asymmetric unit with visible electron density for residues 6 – 96 in each chain (Fig. 3A). Pfc_EnFtn was refined at 2.03 Å resolution and contained three decamers in the asymmetric unit, with visible electron density for residues 2–98 in each chain (Fig. 3B). The overall architecture of both structures mirrors the annular decamer seen in the structure of Rru_EncFtn (Fig. 3C). The electrostatic surfaces of these proteins display similar features to Rru_EncFtn, with negatively charged patches around the circumference that correspond to the exterior metal binding sites seen on Rru_EncFtn, and a negatively charged tunnel at the centre of the decamer corresponding to the interior metal binding site (Fig. S3).

**Figure 3.**
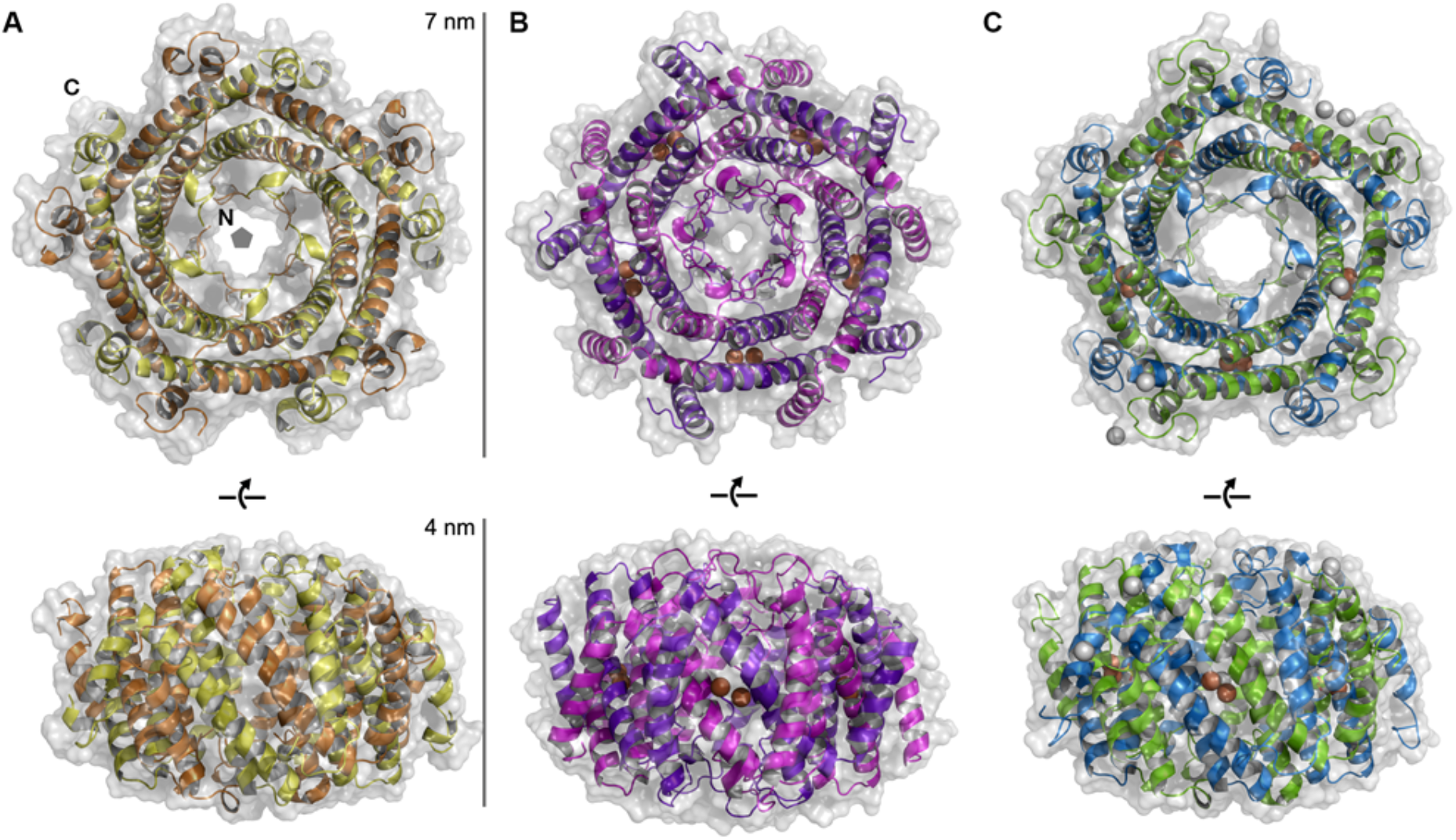
Encapsulated ferritins from *Haliangium ochraceum* and *Pyrococcus furiosus* form annular decamers. The annular decameric architecture of the encapsulated ferritins from *Haliangium ochraceum, A*, and *Pyrococcus furiosus, B*, are shown as transparent solvent accessible surfaces over secondary structure cartoons. The published structure of the *Rhodospirillum rubrum* encapsulated ferritin is shown for comparison, *C* (PDB ID: 5DA5) (1). Bound metal ions are shown as spheres: iron ions are depicted in orange, and calcium ions in grey. The positions of the N- and C-termini of the protein chains and five-fold symmetry axis are highlighted in panel *A*.

**Table 1.** X-ray diffraction data collection and refinement statistics.

|  | Hoch_EncFtn | Pfc_EncFtn* |
| --- | --- | --- |
| **Data collection** |  |  |
| Wavelength (Å) | 1.72 | 1.74 |
| Resolution range (Å) | 47.82 - 2.06<br>(2.131 - 2.06) | 47.19 - 2.03<br>(2.099 - 2.03) |
| Space group | C 1 2 1 | P 1 21 1 |
| Unit cell (Å) | 91.63, 92.65, 119.29 | 99.85, 110.06, 136.27 |
| β(°) | 106.94 | 91.22 |
| Total reflections | 30,0549 (27,242) | 1,226,338 (112,156) |
| Unique reflections | 57,293 (5,570) | 374,905 (36,076) |
| Multiplicity | 5.2 (4.9) | 3.3 (3.0) |
| Completeness (%) | 96.35 (94.08) | 96.95 (95.44) |
| Mean I/sigma(I) | 10.16 (2.37) | 10.48 (4.02) |
| Wilson B-factor (Å<sup>2</sup>) | 31.67 | 22.84 |
| R<sub>merge</sub> | 0.094 (0.596) | 0.080 (0.363) |
| R<sub>meas</sub> | 0.105 (0.667) | 0.095 (0.441) |
| R<sub>pim</sub> | 0.044 (0.292) | 0.052 (0.247) |
| CC1/2 | 0.997 (0.909) | 0.995 (0.902) |
| CC* | 0.999 (0.976) | 0.999 (0.974) |
| Image DOI | 10.5281/zenodo.322743 | 10.5281/zenodo.344797 |
| **Refinement** |  |  |
| Reflections used in refinement | 57,082 (5,563) | 365,581 (36,036) |
| Reflections used for R-free | 2,765 (284) | 17,934 (1697) |
| R<sub>work</sub> | 0.201 (0.295) | 0.175 (0.223) |
| R<sub>free</sub> | 0.245 (0.345) | 0.202 (0.245) |
| CC(work) | 0.964 (0.926) | 0.967 (0.923) |
| CC(free) | 0.936 (0.851) | 0.953 (0.903) |
| Number of non-hydrogen atoms | 7,819 | 25,735 |
| macromolecules | 7,650 | 23,691 |
| ligands | 1 | 30 |
| solvent | 168 | 2,014 |
| Protein residues | 916 | 2,922 |
| RMS(bonds) (Å) | 0.003 | 0.002 |
| RMS(angles) (°) | 0.46 | 0.44 |
| Ramachandran |  |  |
| favoured (%) | 98.77 | 99.90 |
| allowed (%) | 1.23 | 0.10 |
| outliers (%) | 0.00 | 0.00 |
| Rotamer outliers (%) | 0.96 | 0.39 |
| Clashscore | 1.20 | 1.15 |
| Average B-factor (Å<sup>2</sup>) | 44.57 | 28.97 |
| macromolecules | 44.59 | 28.32 |
| ligands | 34.68 | 19.73 |
| solvent | 43.70 | 36.74 |
| PDB ID | 5N5F | 5N5E |

\*Anomalous pairs were counted separately for Pfc\_EncFtn, as this model was refined against the anomalous data. Statistics for the highest-resolution shell are shown in parentheses.

The monomers of Hoch_EncFtn and Pfc_EncFtn superimpose with an RMSD of 1.21 Å over 74 Cα atoms. Hoch_EncFtn superimposes on Rru_EncFtn with an RMSD of 0.47 Å over 91 Cα atoms, while Pfc_EncFtn superimposes with an RMSD of 1 Å over 71 Cα atoms. While the Hoch_EncFtn and Rru_EncFtn structures are almost identical, the Pfc_EncFtn structure presents several key differences to these two proteins. At the N-terminus of Pfc_EncFtn, there is visible electron density from Gly2, whereas the chains of both Hoch_EncFtn and Rru_EncFtn are not visible until residue Gln6 and Ser7 respectively. The additional structured residues in Pfc_EncFtn form an extended loop (Fig. 4), which lies in the central channel of the physiologically active decamer and forms a rigid constriction when compared to the Hoch_EncFtn and Rru_EncFtn structures (Fig 3B). The C-terminal a3 helix of Pfc_EncFtn has two additional turns when compared to the other structures and is shifted by 25° relative to a2 (Fig. 4B/C); this extends its interaction with the neighbouring dimer. (Fig. 3B).

**Figure 4.**
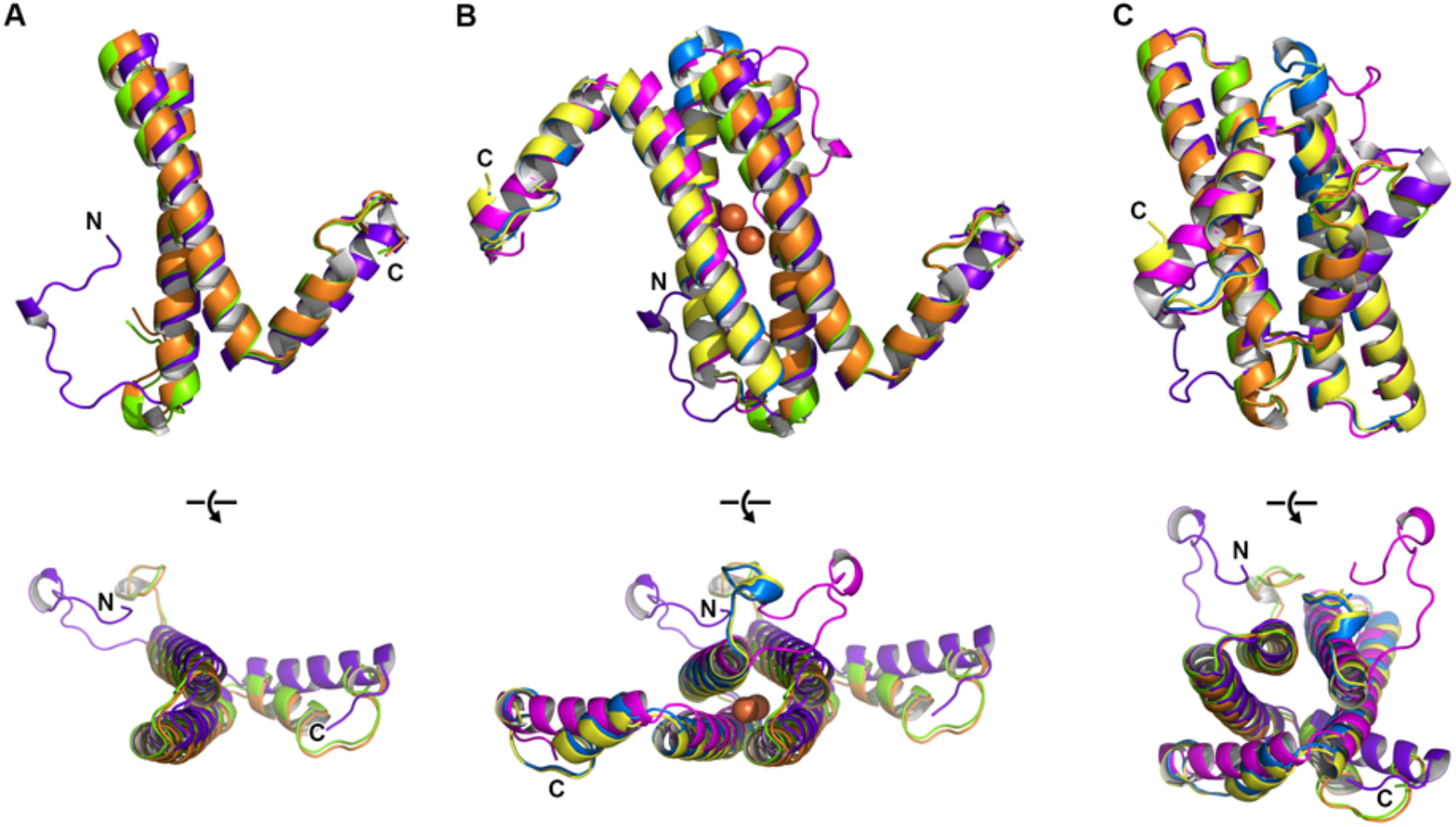
Crystal structures of encapsulated ferritins from *Haliangium ochraceum* and *Pyrococcus furiosus*. Secondary structure cartoon depictions of the structures of the protomers and dimers found in the crystal structures of the Hoch_EncFtn and Pfc_EncFtn encapsulated ferritins. *A*, orthogonal views of the protomers, Hoch_EncFtn shown in orange; Pfc_EncFtn in purple; and Rru_EncFtn in green for comparison (PDB ID: 5DA5) (1). *B*, orthogonal views of the ferroxidase centre dimer. Iron ions present in the Pfc_EncFtn dimer are shown as orange spheres. No metal ions were present in the Hoch_EncFtn structure. Hoch_EncFtn is shown in orange/yellow; Pfc_EncFtn in purple/pink; and Rru_EncFtn in green/blue. *C*, orthogonal views of the non-FOC dimer, depicted as in *B*.

The main interaction surfaces that make up the decameric arrangement in the proteins correspond to the ferroxidase centre dimer (FOC) and the non-ferroxidase dimer (non-FOC) interfaces (Fig. 3B/C). Analysis of the extent of these surfaces with PISA (14) gives a buried surface of 1186 Å^2^ for the Hoch_EncFtn FOC interface, with 8 hydrogen bonds and 16 salt bridges; and 1712 Å^2^ in Pfc_EncFtn, with 14 hydrogen bonds and 6 salt bridges (Table S2). While Hoch_EncFtn buries roughly the same surface area in its FOC interface as Rru_EncFtn FOC (1267 Å^2^), the latter has only 2 hydrogen bonds and 6 salt bridges; the additional stabilisation of this interface in Hoch_EncFtn is likely related to the environmental niche of *Haliangium ochraceum*, as proteins from halophilic organisms tend to have an increased number of salt bridges when compared to those from mesophiles (15). The Pfc_EncFtn FOC interface has fourteen hydrogen bonds and six salt bridges; this significant increase in hydrogen bonding over the Rru_EncFtn protein is likely to be a consequence of the hyperthermophilic nature of *Pyrococcus furiosus*.

The non-FOC interface for Hoch_EncFtn buries 2338 Å^2^, which is similar to the 2468 Å^2^ buried in this region in Rru_EncFtn; however, there are 18 hydrogen bonds compared to 16 in Rru_EncFtn, and 5 salt bridges compared to 16 in Rru_EncFtn (Table S2). These differences are due to a higher proportion of buried hydrophobic residues in the Hoch_EncFtn interface. The non-FOC interface in the Pfc_EncFtn is less extensive than either of the other two structures at 1793 Å^2^, with 19 hydrogen bonds and 19 salt bridges. The extended structured loop at the N-terminus and shifted α3 helix form one hydrogen bond each to the monomer next to their partner in the FOC interface, to bury an additional 586 Å^2^ of surface area. Taken together, these two interfaces bury about the same surface area as the non-FOC interfaces of both Rru_EncFtn and Hoch_EncFtn. The extended interaction surfaces in the Pfc_EncFtn potentially act as a girdle around the decamer, further stabilising it at high temperatures.

The residues in the iron binding site of the FOC interface are conserved in the Hoch_EncFtn and Pfc_EncFtn structures (Fig. 5). The Pfc_EncFtn structure shows clear anomalous difference density in data collected close to the iron edge at 1.74 Å (Fig S4); therefore, a dinuclear iron centre was modelled into the FOC of this structure (Fig. 5). The metal coordination in Pfc_EncFtn is identical to that seen in Rru_EncFtn, with glutamic acid carboxyl oxygen to iron coordination distances of 2.1 Å, and histidine nitrogen to iron distances of 2.2 Å. The electron density in the FOC interface of the Hoch_EncFtn crystal does not contain any peaks for coordinated metal ions (Fig. S4). Given the importance of the histidine residue (His64) for iron coordination (1) and the fact that the protein crystallised in a buffer at pH 5.5, this histidine residue is likely to be around 70 % protonated and would be unable to coordinate iron in this state, thus explaining the absence of iron in this site. In the absence of iron in the Hoch_EncFtn structure, the side-chain of Glu31 is flipped 180° and is within hydrogen bonding distance of Tyr38 from the partner chain, presumably stabilising the apo-form of this interface (Fig. 5A).

**Figure 5.**
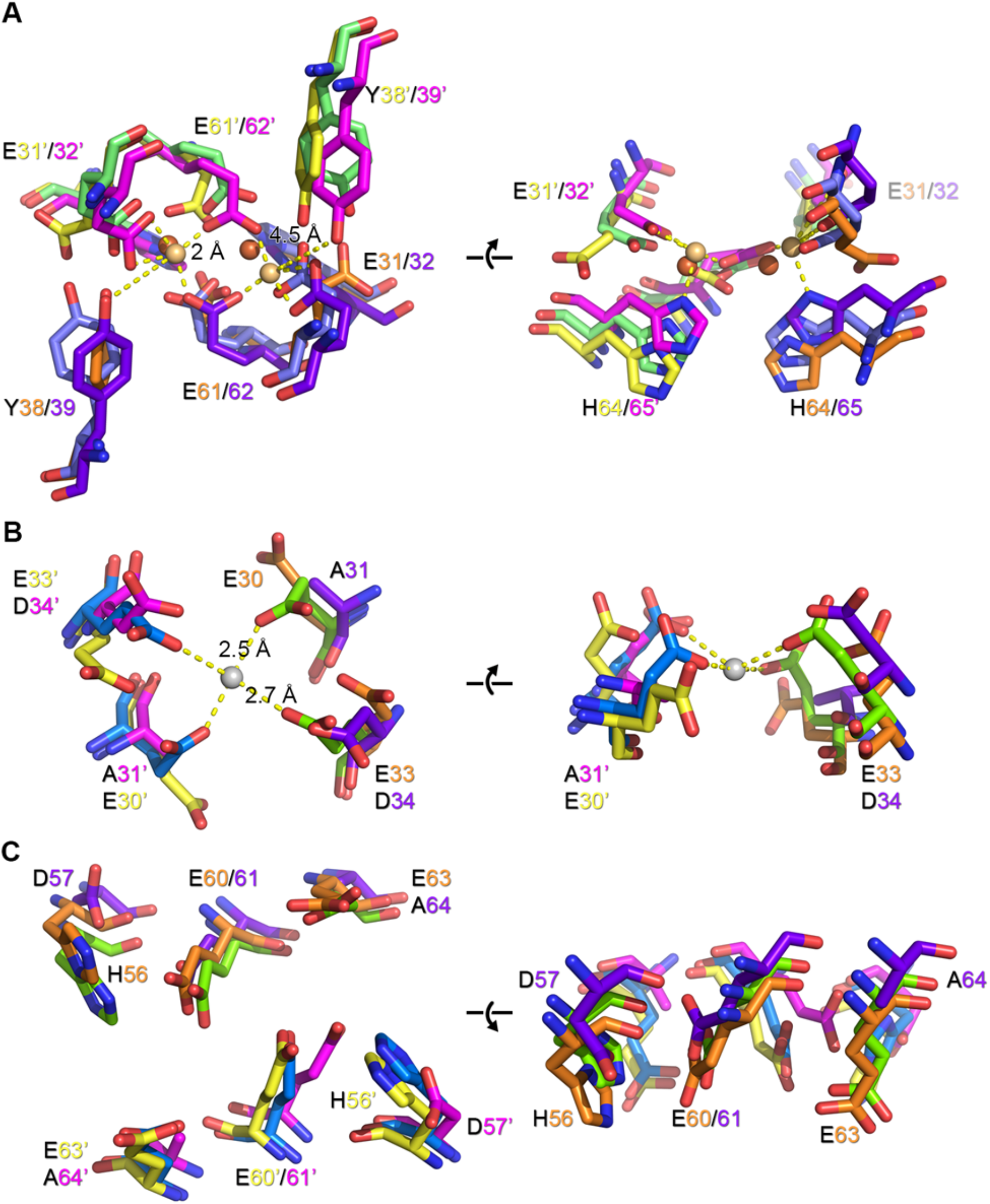
Ferroxidase centre and putative metal binding sites of encapsulated ferritins. Orthogonal views of the ferroxidase centre (*A*) and putative secondary metal binding sites (*B/C*) of Hoch_EncFtn (yellow/orange) and Pfc_EncFtn (pink/purple) compared to Rru_EncFtn (PDB ID: 5DA5) (1) (blue/green). *A*, conserved ferroxidase centre residues are numbered for Hoch_EncFtn and Pfc_EncFtn. Iron ions found in the Pfc_EncFtn FOC are shown as gold spheres with coordination distances to side chain atoms shown. The positions of the iron ions in the Rru_EncFtn structure are shown as orange spheres. *B*, residue conservation in the site occupied by calcium in the Rru_EncFtn structure (grey sphere, with coordination distances). *C*, conserved residues on the outer surface of the encapsulated ferritin surface, located 10 Å from the FOC.

The secondary metal coordination sites seen in the structure of Rru_EncFtn are fully conserved in Hoch_EncFtn and only partially so in Pfc_EncFtn. The metal binding site at the centre of the decameric ring comprises Glu30/Glu33 in Hoch_EncFtn and Ala31/Asp34 in Pfc_EncFtn. Neither structure has any coordinated metal ions in this site (Fig. 5B); The side-chains of the glutamic acid residues are shifted when compared to those in Rru_EncFtn; this could be linked to the absence of a metal ion. The external metal binding site of Hoch_EncFtn is identical to the Rru_EncFtn site, while the Pfc_EncFtn has a glutamic acid in place of a histidine and alanine in place of a glutamic acid in this site.

Given the observation that a decameric form of Hoch_EncFtn was obtained in crystals at pH 5.5, the influence of pH on protein oligomerisation was investigated using native MS. Experiments were performed in pH 5.5 ammonium acetate, and under these acidic conditions MS analysis reveals a substantial increase in the abundance of the decameric species (21+ to 25+) (Fig. S5), similar to the level seen for Pfc_EncFtn and Rru_EncFtn. The lower order oligomerisation states, present at pH 8.0, are significantly reduced in abundance and only dimer (9+ and 10+) and tetramer (14+ and 15+) minor species are observed (Figure S5, green and purple circles respectively). Taken together, our native MS observations suggest that a stable Hoch EncFtn dimer is readily formed irrespective of pH; and under acidic conditions, Hoch_EncFtn dimers favour assembly into the higher order pentamer-of-dimers annular structure which is characteristic of this class of protein.

### Ferroxidase activity

Given the absolute conservation of the FOC residues in Hoch_EncFtn, Pfc_EncFtn and Rru_EncFtn, the ferroxidase activity of the EncFtn homologues was tested to determine if they were indeed active as ferroxidase enzymes. Encapsulated ferritin variants were assayed for their ferroxidase activity by following progress curves for iron oxidation at 315 nm. Each of the proteins displayed similar activity profiles, with some small differences in the shape of the progress curves. Hoch_EncFtn exhibits a higher initial rate than the other homologues (Fig. 6A). Overall these data confirm the ferroxidase activity of the EncFtn family across different species from distinct environmental niches.

**Figure 6.**
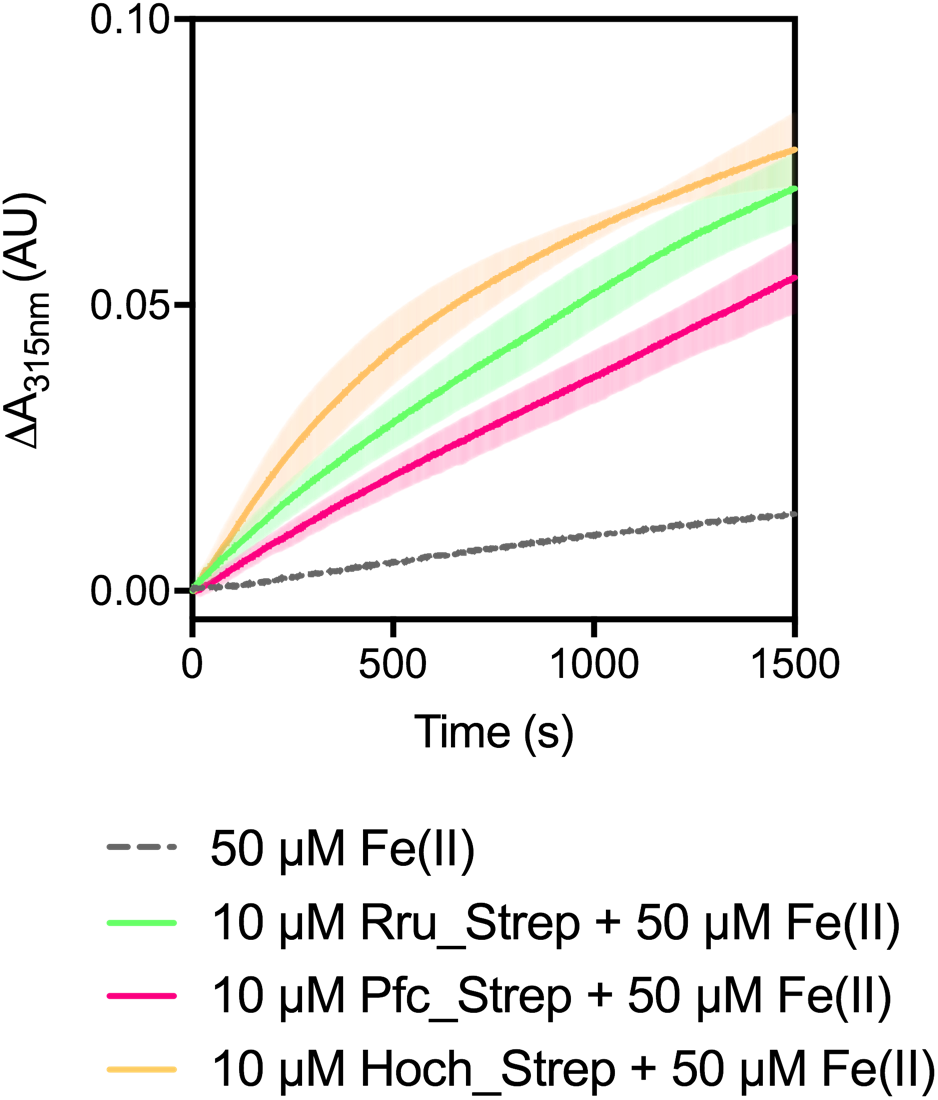
Ferroxidase activity of encapsulated ferritins. Hoch_Strep, Pfc_Strep, and Rru_Strep (10 μM monomer) were incubated with 50 μM FeSO_4_.7H_2_O (10 times molar equivalent Fe^2+^ per FOC) and progress curves of the oxidation of Fe^2+^ to Fe^3+^ was monitored at 315 nm. The background oxidation of iron at 50 μM in enzyme-free control is shown for reference. Solid lines represent the average (n = 3) of technical replicates, shaded areas represent standard deviation from the mean. DOI: 10.6084/m9.figshare.7105685

The metal content of the purified protein fractions used for enzymatic assays was analysed by inductively coupled plasma mass spectrometry (Table S3). Iron levels determined in monomeric Hoch_EncFtn variants and decameric Pfc_Encftn and Rru_EncFtn variants show an iron to protein ratio consistent with 20–80 % occupancy of the ferroxidase centre, assuming all of the iron is located within this site. A ~50 % occupancy of the ferroxidase centre was also observed with the His-tagged EncFtn variants which were used in the mass-spectrometry analysis.

Significant amounts of zinc were copurified with all protein samples in this study, with varying levels detected in the forms with different tags. Zinc is known to strongly inhibit the ferroxidase activity of ferritin family proteins via competitive binding to the ferroxidase centre (16). Given the structural differences between encapsulated ferritins and the classical ferritins, and the differences in secondary metal-binding sites displayed by Pfc_EncFtn, the inhibitory effect of zinc on the EncFtn homologues was tested by performing the ferroxidase assays in the presence of increasing concentrations of zinc. The results show that increasing zinc concentrations lead to a decrease in the slope and final level of ferroxidase progress curves for all of the EncFtn variants, with a maximum inhibition seen with ~ 40–50 μM Zn (Figure S6). These data were fitted using a nonlinear regression with a dose-response model with three parameters, where the response (zero-subtracted end-point absorbance at 315 nm) was the average of three or more replicates per condition (Fig. 7). The calculated IC50 values for zinc inhibition of the three EncFtn variants (Table S4) show significant differences, with the Rru_EncFtn showing the lowest, and Hoch_EncFtn the highest IC50 values. The lower susceptibility of the Hoch_EncFtn to zinc dependent inhibition could be a consequence of a higher level of ferroxidase activity, or structural differences that influence metal binding and discrimination.

**Figure 7.**
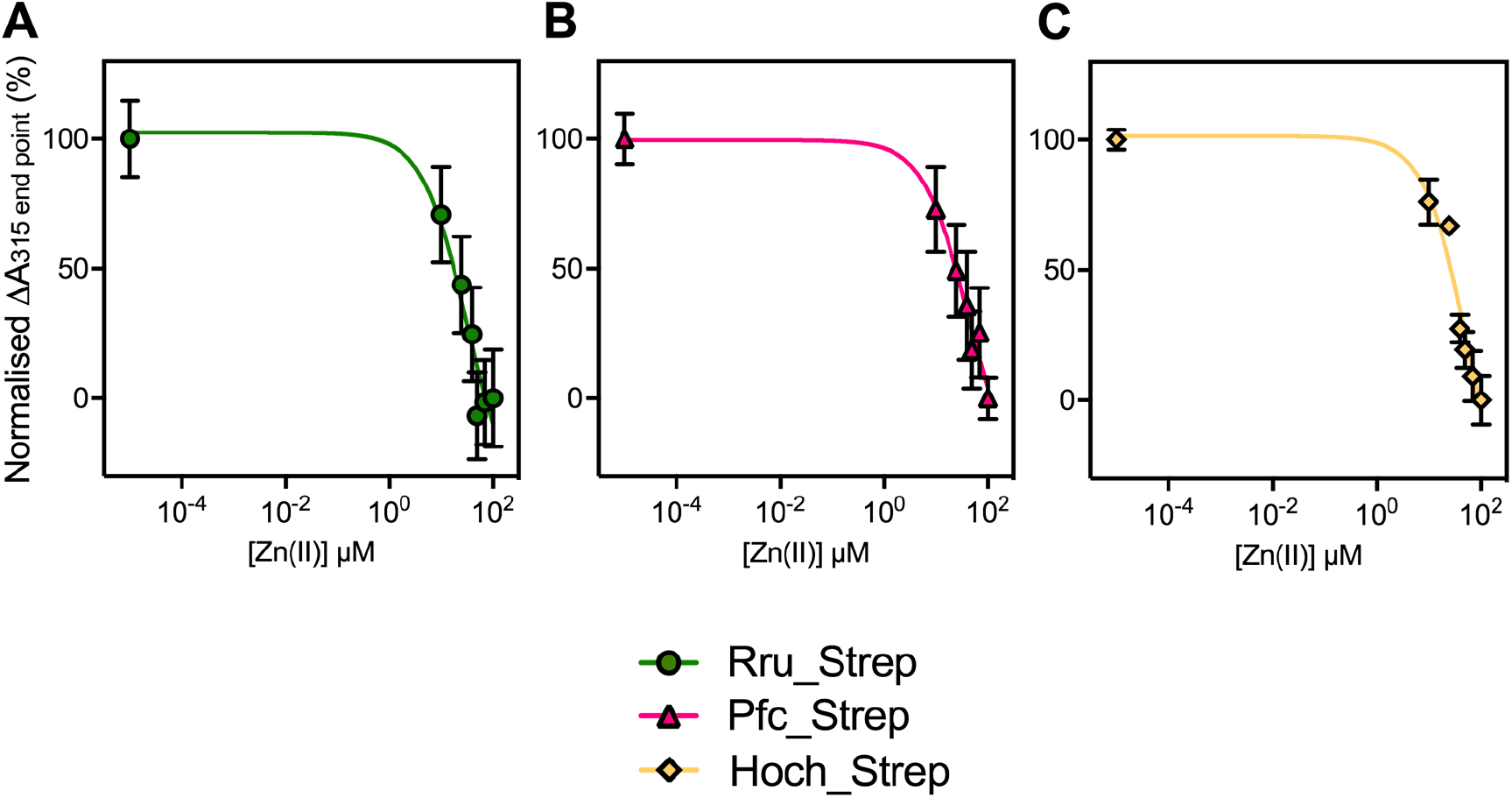
The ferroxidase activity of encapsulated ferritins is inhibited by zinc. *A*/*B*/*C* Non-linear fit of ferroxidase activities of Strep-tagged proteins inhibited by varying concentrations of Zn(II) by GraphPad software using a Dose-Response (log(inhibitor) vs. response (three parameters) equation. Response used in this analysis is the average of three or more technical replicates per condition. Data shown have been recorded after 1500 seconds from the start of the assay. IC50 and logIC50 have been calculated for each protein (Table S4). DOI: 10.6084/m9.figshare.7105685

## Discussion

The recently published structural and functional analysis of the *R. rubrum* encapsulated ferritin system presented a new functional organisation for iron mineralisation by ferritin-like proteins, with the four-helix bundle ferritin-like fold formed through the interaction of EncFtn subunits to form a functional ferroxidase centre (1). The open decameric structure of the EncFtn protein is not competent to store iron in mineral form; instead, the interaction with, and sequestration within, the encapsulin nanocage provides a functional and high-capacity iron storage system (3, 4). To understand whether the structural organisation of the EncFtn family is conserved across microorganisms with different environmental niches and distinct encapsulin geometries, we determined the structure and biochemical properties of EncFtn proteins from *H. ochraceum* and *P. furiosus*.

*P. furiosus* is a hyperthermophilic Euryarchaeota and its encapsulin was the first to be structurally characterised; however, it was initially mis-annotated as a non-functional virus-like particle (13). The *P. furiosus* encapsulin forms a contiguous polypeptide chain with an N-terminal EncFtn domain appended to it and assembles into a 180 subunit T=3 capsid. This archaeal arrangement is distinct to the bacterial EncFtn/encapsulin systems, which are usually encoded in a two-gene operon with the EncFtn upstream of the encapsulin, and with a short C-terminal encapsulation sequence peptide appended to the encoded EncFtn protein (5, 10). It is not clear whether the genomic arrangement of the bacterial EncFtn/encapsulin systems arose separately, or by horizontal gene transfer from archaea followed by mutation of the single-reading frame into two, along with the mutation of the T=3 form of the encapsulin to the T=1 form found in the bacterial EncFtn encapsulins.

The published crystal structure of the *P. furiosus* encapsulin lacks any electron density for the EncFtn domain, implying that this domain is mobile within the encapsulin nanocage even though it is contiguous with and tethered to the encapsulin protein (13). Our analysis focused on the isolated *P. furiosus* EncFtn domain, which forms a decamer in solution, the gas phase, and in crystals. This domain is partially resistant to thermal and SDS induced denaturation, which is in accord with the extreme temperatures endured by the host organism in its volcanic niche.

The current model for the organisation of bacterial EncFtn proteins within encapsulin nanocages places the decamers at the pentameric vertices of the T=1 capsid (1, 4) with the encapsulation sequences of five subunits captured by clefts on the interior surface of the penton of the encapsulin shell. The encapsulation sequences of the other five subunits in the decamer are disengaged and free within the interior of the capsid. No evidence is available on the strength of the interaction between the capsid and the EncFtn protein; however, the presence of clear electron density for some of the encapsulation sequence in the *T. maritima* encapsulin structure implies that the interaction is relatively stable, and this is enhanced by an avidity mechanism with multiple encapsulation sequences engaged by subunits at the pentameric vertex (4). In the archaeal systems the EncFtn and encapsulin domains are a contiguous polypeptide, with the EncFtn domain tethered to the interior of the capsid. The T=3 geometry of the *P. furiosus* encapsulin has 180 subunits, with 12 penton units and 20 hexons. Given the structural and biochemical conservation of the decameric EncFtn protein, it is likely that this arrangement is found within the T=3 as well as T=1 capsids. It is not clear how the decamer could be formed from tethered subunits unless the EncFtn domains found in hexons have enough flexibility in the linker between domains to engage with partners in adjacent pentons. With full engagement at the pentons, this would leave 60 ‘free’ EncFtn domains within the capsid. This observation, coupled with our solution and gas phase experiments, indicates that the quaternary structure of EncFtn proteins is dynamic and that they can exist in equilibrium between monomers/dimers and higher order multiples of dimers.

The dynamic nature of EncFtn proteins is highlighted in our solution and gas phase analyses of the *H. ochraceum* EncFtn. Despite high sequence identity and the conservation of key residues between it and the *R. rubrum* EncFtn, this protein is less prone to multimerisation in solution and displays a greater range of oligomerisation intermediates than both the *R. rubrum* and *P. furiosus* EncFtn proteins in the gas phase. The presence of the conserved EncFtn decamer quaternary structure in the crystal highlights the conservation of this architecture. The absence of metal ions in the crystal structure, which was formed at pH 5.5, indicated that the quaternary structure can be induced by both metal binding, as was shown in solution for the *R. rubrum* EncFtn (1), and changes in pH, presumably through the protonation of the conserved histidine at low pH and the formation of stabilising hydrogen bonds by residues normally involved in the formation of the FOC.

Analysis of the structure and sequence of the *H. ochraceum* EncFtn indicates that the dimerisation interfaces are less extensive than the *R. rubrum* EncFtn interfaces and are discontinuous. This leads to lower binding energies for both of the interfaces when compared to both *R. rubrum* and *P. furiosus*, and this could explain the differences in stability seen in solution and the gas phase.

We show here that the three proteins in this study exhibit comparable ferroxidase activities. This indicates that they were purified in a functional state. Previous reports show that ferritins are susceptible to inhibition by Zn(II) ions (16), which is also the case for the EncFtn family as demonstrated in this study. In the experimental conditions used in our study, the *H. ochraceum* EncFtn was slightly more active than the other proteins and less susceptible to inhibition by zinc. This may be a consequence of the more dynamic nature of this homologue, as seen in solution and the gas phase. The precise mode of catalysis for ferritins is a subject of some debate and key questions as to whether the iron ions engaged within the ferroxidase centre are labile, or act as a stable prosthetic centre; and whether the two iron ions move in concert are the subject of some controversies in the field (17, 18). Our study does not aim to address these controversies, but we do note that the location of the conserved FOC and secondary metal binding sites at a dimer interface and the dynamic nature of EncFtn oligomerisation may indicate a distinct mechanism of iron oxidation and transfer to minerals when compared to the classical ferritins, thus adding a new level of complexity and debate to the field. It must also be noted that the activity of EncFtn proteins occurs within the privileged environment of an encapsulin shell, which adds an additional level of complexity to the study of their catalysis.

The Pfc_EncFtn protein has the lowest enzymatic activity in the set. Analysis of the sequence alignment (Fig. S1) highlights the absence of conserved residues found in the putative metal ion entry site found in the other EncFtn homologues. We propose that the acidic residues found in this location attract iron ions and channel them to the FOC. Their absence in Pfc_EncFtn implies that metal ions reach its FOC simply by diffusion, resulting in a slower Fe(II)/Fe(III) turn-over and hence a reduction in enzyme activity. It would be interesting to apply a mutagenesis approach to further explore the role of these proposed entry site residues on the ferroxidase activity.

Further study will shed light on the role of the different dimerization interfaces on the stability of the EncFtn decamer; and the role of the secondary metal binding sites on catalysis, metal selectivity, and catalytic inhibition by competing metal ions.

## Experimental Procedures

### Cloning expression and purification of EncFtn and EncFtn homologues

Recombinant encapsulated ferritin from *Rhodospirillum rubrum* was produced as described previously (1). DNA fragments encoding truncated versions of the encapsulated ferritins from *Haliangium ochraceum* (Hoch_3836_1–98_) and *Pyrococcus furiosus* (Pfc_01575_1–99_) were produced as double-stranded gBlocks (IDT) and codon optimised for expression in *E. coli*, with restriction endonuclease sites for insertion into pET-28a (Pfc_05157_1–99_) or a modified pET-28 vector with CIDAR MoClo (19) Golden Gate cloning sites (Hoch_3836_1–98_). Untagged and hexa-histidine tagged variants were produced for both proteins in this way. A StrepII-tagged variant of each protein was produced by assembly of the EncFtn gBlock into a CIDAR MoClo destination vector with a custom T7 promoter and StrepII-tag terminator part. Sequences of primers and gBlocks used in this study are shown in Tables S5 and S6 respectively, and the protein sequences of expressed proteins are shown in Table S7. The expression plasmids were transformed into *E. coli* BL21(DE3) cells and a single colony was grown overnight at 37 °C in 100 ml LB medium, supplemented with 35 μg/ml kanamycin, with shaking at 180 rpm. The cells were sub-cultured into 1 litre of LB or M9 minimal medium, grown until OD_600_=0.6, and protein production was induced with 1 mM IPTG, the temperature was reduced to 18 °C and cells were incubated for a further 18 hours.

Cells were harvested by centrifugation at 4,000 × *g* and washed with PBS. Cell-free extract was produced by resuspending cells in 10 x v/w Buffer A (50 mM Tris-HCl, pH 8.0) and sonicated on ice for 5 minutes with 30 s on/off cycle at 60 watts power output. The lysate was cleared by centrifugation at 35,000 × *g* and filtered with a 0.45 μm syringe filter. Untagged recombinant proteins were purified from cell-free extract by anion exchange using HiPrep Q sepharose fast-flow 16/10 column (GE Healthcare) equilibrated with Buffer A. Cell free extract was applied to the column and unbound proteins were washed off with 10 column volumes of Buffer A. Proteins were eluted with a linear gradient of Buffer B (50 mM Tris-HCl, pH 8.0, 1 M NaCl) over 20 column volumes and fractions collected.

Hexa-histidine tagged proteins were purified by resuspending cells in 10 x (v/w) Buffer HisA (50 mM Tris-HCl, pH 8.0, 500 mM NaCl, 50 mM imidazole) before sonication and clarification as above. Clarified cell lysate was applied to a 5 ml HisTrap FF column (GE Healthcare) and unbound proteins were washed off with 10 column volumes of Buffer HisA. A step-gradient of 50 % and 100 % Buffer HisB (50 mM Tris-HCl, pH 8.0, 500 mM NaCl, 500 mM imidazole) was used to elute His-tagged proteins.

Fractions of Q sepharose or His-trap eluent containing the protein of interest, as identified by 15 % (w/v) SDS-PAGE, were subjected to size-exclusion chromatography using an S200 16/60 column equilibrated with Buffer GF (50 mM Tris-HCl, pH 8.0, 150 mM NaCl).

StrepII-tagged proteins were purified by suspending cells in 10 x (v/w) Buffer W (100 mM Tris pH 8.0, 150 mM NaCl) before sonication and clarification as described above. Cell lysate was applied to Strep-Trap HP column (GE Healthcare), prepared as suggested by the manufacturer, and unbound proteins were washed off by applying 5 column volumes of Buffer W. Strep-tagged proteins were eluted by Buffer E (100 mM Tris pH 8.0, 150 mM NaCl, 2.5 mM desthiobiotin). Eluted protein was buffer-exchanged to Buffer GF by centrifugal concentrator Vivaspin Turbo (Sartorius, 10 kDa MWCO) to remove desthiobiotin.

### Ferroxidase assays

Ferroxidase activity of the enzymes was tested by following the formation of Fe(III) species by UV-visible spectroscopy as previously described (1). Oxygen-free aliquots of Fe(II) (1 mM) were prepared by dissolving FeSO_4_.7H_2_O in 0.1 % (v/v) HCl under anaerobic conditions. Purified proteins (decameric fraction) were diluted anaerobically to the final concentration of 10 μM monomer in Buffer H (10 mM HEPES pH 8.0, 150 mM NaCl). Protein and Fe(II) samples were added to a quartz cuvette (Hellma) under aerobic conditions at the final concentration of 10 and 50 μM, respectively. Absorbance at 315 nm was monitored every second over 1500 s using a UV-visible spectrophotometer (Perkin Elmer Lambda 35) using the provided TimeDrive software. A negative control was performed by monitoring the progress curve at A_315_ of Fe(II) salt sample in the absence of protein. Data presented here are the mean of three technical replicates of time zero-subtracted progress curves with standard deviations calculated from the mean.

### Zinc inhibition of ferroxidase activity

In order to test enzyme selectivity toward Fe(II), the ferroxidase assay was carried out as previously described (1), in the presence of FeSO_4_.7H_2_O (50 μM) and various concentrations of ZnSO_4_.2H_2_O. The A_315_ nm progress curve of protein mixed with the highest concentration of Zn(II) used in the assay (100 μM) was also monitored as a negative control.

### Protein Crystallography

Hoch_EncFtn and Pfc_EncFtn were concentrated to 10 mg/ml using a 10 kDa MWCO centrifugal concentrator (Vivaspin) and subjected to sitting drop vapour diffusion crystallisation using 70 μl well solution and drops of 100 nl protein plus 100 nl well solution. Crystals of Hoch_EncFtn grew in well solution containing 0.2 M NaCl, 0.1 M Bis-Tris, pH 5.5, 20 % (w/v) PEG 3350; and crystals of Pfc_EncFtn were found in a condition containing 0.2 M LiSO_4_.H_2_O, 20 % (w/v) PEG 3350. Crystals were harvested using a LithoLoop (Molecular Dimensions Limited), transferred to a cryoprotection solution of well solution supplemented with 20 % (v/v) PEG 200 and flash cooled in liquid nitrogen. Diffraction data were collected at Diamond Light Source at 100 K using a Pilatus 6M detector. Images were integrated and scaled using XDS (20); the correct crystallographic symmetry group was confirmed with Pointless (21) and reflections were merged with Aimless (22). The data were phased by molecular placement using Phaser, with a decamer of Rru_EncFtn (PDB ID: 5DA5) used as the search model; this was edited to match the target sequence using CHAINSAW (23). The crystallographic models were rebuilt using phenix.autobuild (24) and subsequently refined using phenix.refine (25) with cycles of manual model building in Coot (26). The model quality was assessed using MolProbity (27). All structural figures were generated using PyMOL (www.pymol.org). X-ray data collection and refinement statistics are shown in Table 1.

### Sequence alignment and depiction

The protein sequences for Hoch_3836 and the encapsulated ferritin domain of Pfc_05175 were aligned against Rru_A0973 using Clustal Omega (28) and rendered using ESPript 3.0 (29).

### Mass spectrometry

All mass spectrometry (MS) experiments were performed on a Synapt G2 ion-mobility equipped Q-ToF instrument (Waters Corp., Manchester, UK). LC-MS experiments were performed using an Acquity UPLC equipped with a reverse phase C4 Aeris Widepore 50 × 2.1 mm HPLC column (Phenomenex, CA, USA) and a gradient of 5– 95% acetonitrile (0.1% formic acid) over 10 minutes was employed. For LC-MS, samples were typically analysed at 5 μM, and data analysis was performed using MassLynx v4.1 and MaxEnt deconvolution. For native MS analysis, all protein samples were buffer exchanged into 100 mM ammonium acetate (pH 8.0, or pH 5.0) using Micro Biospin Chromatography Columns (Bio-Rad, UK) prior to analysis and the resulting protein samples were analysed at a typical final concentration of ~5 μM (oligomer concentration). For native MS ionisation, nano-ESI was employed using a nanomate nanoelectrospray infusion robot (Advion Biosciences, Ithaca, NY). Instrument parameters were tuned to preserve non-covalent protein complexes and were consistent for the analysis of all protein homologues. After the native MS optimisation, parameters were: nanoelectrospray voltage 1.60 kV; sample cone 100 V; extractor cone 0 V; trap collision voltage 4 V; source temperature 60 °C; and source backing pressure 6.0 mbar.

### ICP-MS

Inductively coupled plasma mass spectrometry experiments were performed on samples of Rru_His, Hoch_His, and Pfc_His from size-exclusion chromatography experiments and Rru_Strep, Hoch_Strep, and Pfc_Strep from the affinity chromatography purification step as described previously (1).

### Data Availability

Mass spectrometry data presented in this study is available at Edinburgh Datashare (http://dx.doi.org/10.7488/ds/2443).

Protein structure models and electron density data are available at the PDB (Hoch_EncFtn, 5N5F; Pfc_EncFtn, 5N5E). Diffraction images are available at Zenodo (Hoch_EncFtn, 10.5281/zenodo.322743; Pfc_EncFtn, 10.5281/zenodo.344797). Data for ferroxidase assays are available at FigShare.

## Acknowledgements

This work was supported a Royal Society Research Grant awarded to JMW (RG130585) and a BBSRC New Investigator Grant to JMW and DJC (BB/N005570/1). JMW is funded by Newcastle University. DJC and JR are funded by the University of Edinburgh. JR is funded by a BBSRC EastBio DTP studentship (BB/M010996/1). DH was funded by the China Scholarship Council. LRT was funded by a BBSRC EastBIO DTP Studentship. KJW and ET are funded by the Wellcome Trust and Royal Society through a Sir Henry Dale Fellowship awarded to KJW (098375/Z/12/Z). We would like to thank the staff on the Macromolecular Crystallography beamlines at Diamond Light Source for their assistance with data collection. We would like to thank Prof Dominic Campopiano and Dr Elisabeth Lowe for their critical reading of this manuscript and helpful discussions.

## Author Contributions

DH, JMW and DC conceived the study. DH produced the native and hexa-histidine tagged constructs used in the study. LRT produced the DNA parts and assembled the StrepII-tagged variants of the EncFtn proteins. DH, CP, JR, ZM and JMW produced recombinant proteins used in the study. DH and JMW determined and refined the crystal structures. CP performed biochemical assays shown in Figures 6/7/S6. JR and DC performed mass spectrometry experiments shown in Figures 2 and S5. CLM provided technical assistance and contributed to data collection for MS experiments. KJW and ET performed ICP-MS experiments shown in Table S3. All authors contributed to data analysis and preparation of the manuscript.

## References

1. He, D., Hughes, S., Vanden-Hehir, S., Georgiev, A., Altenbach, K., Tarrant, E., Mackay, C. L. L., Waldron, K. J. K. J., Clarke, D. J. D. J., and Marles-Wright, J. (2016) Structural characterization of encapsulated ferritin provides insight into iron storage in bacterial nanocompartments. Elife. 5, e18972

2. Andrews, S. C. (2010) The Ferritin-like superfamily: Evolution of the biological iron storeman from a rubrerythrin-like ancestor. Biochim. Biophys. Acta. 1800, 691–705

3. McHugh, C. A., Fontana, J., Nemecek, D., Cheng, N., Aksyuk, A. A., Heymann, J. B., Winkler, D. C., Lam, A. S., Wall, J. S., Steven, A. C., and Hoiczyk, E. (2014) A virus capsid like nanocompartment that stores iron and protects bacteria from oxidative stress. EMBO J. 33, 1896–1911

4. Sutter, M., Boehringer, D., Gutmann, S., Günther, S., Prangishvili, D., Loessner, M. J., Stetter, K. O., Weber-Ban, E., and Ban, N. (2008) Structural basis of enzyme encapsulation into a bacterial nanocompartment. Nat. Struct. Mol. Biol. 15, 939–947

5. Giessen, T. W., and Silver, P. A. (2017) Widespread distribution of encapsulin nanocompartments reveals functional diversity. Nat. Microbiol. 2, 17029

6. Crow, A., Lawson, T. L., Lewin, A., Moore, G. R., and Le Brun, N. E. (2009) Structural basis for iron mineralization by bacterioferritin. J. Am. Chem. Soc. 131, 6808–13

7. Bakker, G. R., and Boyer, R. F. (1986) Iron incorporation into apoferritin. The role of apoferritin as a ferroxidase. J. Biol. Chem. 261, 13182–13185

8. Bradley, J. M., Moore, G. R., and Le Brun, N. E. (2014) Mechanisms of iron mineralization in ferritins: one size does not fit all. J. Biol. Inorg. Chem. 19, 775–85

9. Chasteen, N. D., and Harrison, P. M. (1999) Mineralization in ferritin: an efficient means of iron storage. J. Struct. Biol. 126, 182–94

10. Sutter, M., Boehringer, D., Gutmann, S., Günther, S., Prangishvili, D., Loessner, M. J., Stetter, K. O., Weber-Ban, E., and Ban, N. (2008) Structural basis of enzyme encapsulation into a bacterial nanocompartment. Nat. Struct. Mol. Biol. 15, 939–947

11. Snijder, J., Kononova, O., Barbu, I. M., Uetrecht, C., Rurup, W. F., Burnley, R. J., Koay, M. S. T., Cornelissen, J. J. L. M., Roos, W. H., Barsegov, V., Wuite, G. J. L., and Heck, A. J. R. (2016) Assembly and Mechanical Properties of the Cargo-Free and Cargo-Loaded Bacterial Nanocompartment Encapsulin. Biomacromolecules. 17, 2522–9

12. Rurup, W. F., Snijder, J., Koay, M. S. T., Heck, A. J. R., and Cornelissen, J. J. L. M. (2014) Self-sorting of foreign proteins in a bacterial nanocompartment. J. Am. Chem. Soc. 136, 3828–32

13. Akita, F., Chong, K. T., Tanaka, H., Yamashita, E., Miyazaki, N., Nakaishi, Y., Suzuki, M., Namba, K., Ono, Y., Tsukihara, T., and Nakagawa, A. (2007) The crystal structure of a virus-like particle from the hyperthermophilic archaeon Pyrococcus furiosus provides insight into the evolution of viruses. J. Mol. Biol. 368, 1469–83

14. Krissinel, E., and Henrick, K. (2007) Inference of macromolecular assemblies from crystalline state. J. Mol. Biol. 372, 774–97

15. DasSarma, S., and DasSarma, P. (2015) Halophiles and their enzymes: negativity put to good use. Curr. Opin. Microbiol. 25, 120–126

16. Pfaffen, S., Abdulqadir, R., Le Brun, N. E., and Murphy, M. E. P. (2013) Mechanism of ferrous iron binding and oxidation by ferritin from a pennate diatom. J. Biol. Chem. 288, 14917–25

17. Ebrahimi, K., Eckhard, B., Hagedoorn, P., and Hagen, W. (2012) The catalytic center of ferritin regulates iron storage via Fe (II)-Fe (III) displacement. Nat. Chem. Biol. 8, 941–948

18. Hagen, W. R., Hagedoorn, P.-L., and Honarmand Ebrahimi, K. (2017) The workings of ferritin: a crossroad of opinions. Metallomics. 9, 595–605

19. Iverson, S. V., Haddock, T. L., Beal, J., and Densmore, D. M. (2016) CIDAR MoClo: Improved MoClo Assembly Standard and New E. coli Part Library Enable Rapid Combinatorial Design for Synthetic and Traditional Biology. ACS Synth. Biol. 5, 99–103

20. Kabsch, W. (2010) Integration, scaling, space-group assignment and post-refinement. Acta Crystallogr. Sect. D Biol. Crystallogr. 66, 133–144

21. Evans, P. R. (2011) An introduction to data reduction: space-group determination, scaling and intensity statistics. Acta Crystallogr. D. Biol. Crystallogr. 67, 282–92

22. Evans, P. (2006) Scaling and assessment of data quality. in Acta Crystallographica Section D: Biological Crystallography, pp. 72–82, International Union of Crystallography, 62, 72–82

23. Stein, N. (2008) CHAINSAW: A program for mutating pdb files used as templates in molecular replacement. J. Appl. Crystallogr. 41, 641–643

24. Terwilliger, T. C., Grosse-Kunstleve, R. W., Afonine, P. V., Moriarty, N. W., Zwart, P. H., Hung, L. W., Read, R. J., and Adams, P. D. (2007) Iterative model building, structure refinement and density modification with the PHENIX AutoBuild wizard. Acta Crystallogr. Sect. D Biol. Crystallogr. 64, 61–69

25. Afonine, P. V., Grosse-Kunstleve, R. W., Echols, N., Headd, J. J., Moriarty, N. W., Mustyakimov, M., Terwilliger, T. C., Urzhumtsev, A., Zwart, P. H., and Adams, P. D. (2012) Towards automated crystallographic structure refinement with phenix.refine. Acta Crystallogr. Sect. D Biol. Crystallogr. 68, 352–367

26. Emsley, P., Lohkamp, B., Scott, W. G., and Cowtan, K. (2010) Features and development of Coot. Acta Crystallogr. D. Biol. Crystallogr. 66, 486–501

27. Chen, V. B., Arendall, W. B., Headd, J. J., Keedy, D. A., Immormino, R. M., Kapral, G. J., Murray, L. W., Richardson, J. S., and Richardson, D. C. (2010) MolProbity: all-atom structure validation for macromolecular crystallography. Acta Crystallogr. D. Biol. Crystallogr. 66, 12–21

28. Sievers, F., and Higgins, D. G. (2014) Clustal Omega, accurate alignment of very large numbers of sequences. Methods Mol. Biol. 1079, 105–16

29. Gouet, P., Robert, X., and Courcelle, E. (2003) ESPript/ENDscript: Extracting and rendering sequence and 3D information from atomic structures of proteins. Nucleic Acids Res. 31, 3320–3

